# *DONSON*, a gene responsible for microcephalic primordial dwarfism, ensures proper centriole duplication cycle by maintaining centriole engagement during interphase

**DOI:** 10.1101/2020.05.10.086777

**Authors:** Kyohei Matsuhashi, Koki Watanabe, Kei K. Ito, Takumi Chinen, Shoji Hata, Grant S. Stewart, Daiju Kitagawa

## Abstract

Microcephalic primordial dwarfism (MPD) is a genetic disorder characterized by short stature and microcephaly. MPD-related genes are known to regulate centrosome biogenesis, DNA replication or the DNA damage response. Although some of the MPD-related proteins that are implicated in DNA replication localize to centrosomes, how these proteins affect centrosome biogenesis remains mostly elusive. Here, we revisit the potential function of these DNA replication mediators in human centrosome biogenesis. Among these proteins, depletion of DONSON leads to excessive number of centrosomes in interphase, caused by precocious centriole disengagement. Such disengaged centrioles are converted to centrosomes, followed by centriole reduplication during interphase. These extra centrosomes lead to abnormal spindle formation and chromosome segregation errors. Importantly, similar defects are observed in MPD patients’ cells with *DONSON* mutations, suggesting a possible cause of the disease. Overall, these results indicate that DONSON is involved in regulating the centriole duplication cycle by ensuring the maintenance of centriole engagement during interphase.

## Introduction

Centrosomes are conserved non-membrane organelles that function as microtubule organizing centers (MTOCs) in most animal cells. Centrosomes play an important role in regulating cell division, cell signaling, motility and many other biological processes (Nigg & Holland, 2018). Each centrosome consists of two centrioles (mother and daughter centrioles) and a surrounding protein matrix known as pericentriolar material (PCM). Centriole duplication starts in the late G1 phase with the assembly of the cartwheel structure at the vicinity of the mother centriole, which is followed by centriole elongation during the S phase. Each daughter centriole is tightly engaged with the mother centriole (centriole engagement) until late mitosis (Loncarek, Hergert, Magidson, & Khodjakov, 2008; Tsou et al., 2009; Tsou & Stearns, 2006). Centriole disengagement occurs in late mitosis, which is promoted by Polo-like kinase 1 (Plk1) and the protease separase (Tsou et al., 2009). Thereafter, the mother centriole is licensed to acquire the ability to recruit PCM and to duplicate a new centriole in the next cell cycle (Nigg & Holland, 2018). Thus, centriole engagement can serve as one of the mechanisms that regulate the centriole number by preventing reduplication of mother centrioles within the same cell cycle (Tsou & Stearns, 2006). The number of centrioles must be tightly regulated to ensure proper spindle formation and chromosome segregation. Aberrations in the expression or function of centriole components may cause abnormal centrosome number and lead to a number of diseases such as cancer and neurodevelopmental disorders (Nigg & Holland, 2018).

Microcephalic primordial dwarfism (MPD) is a term used to describe a diverse group of clinically related disorders, which are characterized by the presence of moderate to severe microcephaly and intrauterine and postnatal growth restriction. It has been proposed that the abnormal neurodevelopment exhibited by the affected patients is often caused by disrupting the cell cycle of neural progenitor cells (NPCs) (Jayaraman, Bae, & Walsh, 2018). Since centrosomal proteins, such as CENPJ, Cep63, and Cep152 (Al-Dosari, Shaheen, Colak, & Alkuraya, 2010; Kalay et al., 2011; Sir et al., 2011) and pericentrin (PCNT, a major PCM component) (Rauch et al., 2008), have all been identified as causative genes for MPD, this suggests that aberrations in the number of centrosomes and abnormal mitotic progression are a major underlying cause of human disease associated with NPC disruption (Nigg & Holland, 2018). In addition to genes involved in centrosome biogenesis, genes involved in regulating the DNA damage response and DNA replication have also been reported to be responsible for causing MPD (Klingseisen & Jackson, 2011). However, despite their identification, it remains poorly understood whether mutations in the MPD genes associated with defects in DNA replication and/or the DNA damage response can also compromise centrosome biogenesis and reduce the pool of NPCs in human brain development.

Interestingly, it has been shown that several DNA replication components linked with MPD, such as Cdc6, Orc1 and GMNN, localize not only in the nucleus, but also at centrosomes, and depletion of these proteins alters centrosome number (Hemerly, Prasanth, Siddiqui, & Stillman, 2009; Kim, Lee, Kim, & Hwang, 2017; Knockleby & Lee, 2010; Lu, Lan, Zhang, Jiang, & Zhang, 2009; Xu et al., 2017). Downstream neighbor of SON (DONSON) is a gene recently identified to be involved in MPD (Evrony et al., 2017; Reynolds et al., 2017). A recent study reported that DONSON is a replisome component that protects stalled or damaged replication forks and maintains genome stability (Reynolds et al., 2017). In addition to regulating the replication fork progression, DONSON is also reported to localize at centrosomes and ensure proper mitotic spindle formation (Fuchs et al., 2010). However, whether DONSON is involved in centrosome biogenesis remains unclear.

In this study, we set up a small siRNA-based screen to explore the potential function of the MPD genes, which have been implicated in DNA replication, in regulating centrosome biogenesis. From this screen, we identified that depletion of DONSON causes precocious centriole disengagement during interphase. Notably, these precociously disengaged centrioles are converted to centrosomes, which are able to duplicate and organize PCM. As a consequence of this, during mitosis, all the disengaged centrioles acquire MTOC activity, which results in multi-polar or pseudo-bipolar spindle formation and chromosome segregation errors. Consistent with DONSON depletion using siRNA, we also find that the cells derived from the patients with hypomorphic *DONSON* mutations also exhibit similar defects, such as precocious centriole disengagement in interphase and abnormal spindle formation in mitosis. Overall, these findings lead us to propose that DONSON not only regulates DNA replication but also the centriole duplication cycle by ensuring timely centriole disengagement in mitosis.

## Results

### DONSON is involved in the regulation of centrosome number

Microcephalic primordial dwarfism (MPD) is a group of genetic disorders characterized by microcephaly with restricted intrauterine growth (Jayaraman et al., 2018). The genes known to be mutated in MPD are mostly classified as either encoding components of the centrosome, the DNA replication machinery or regulators of the DNA damage response (Fig. 1A) (Jayaraman et al., 2018). It has been shown that some of the proteins required for DNA replication localize not only in the nucleus but also to centrosomes, suggesting a potential role for these proteins in regulating centrosome duplication and/or maturation (Hemerly et al., 2009; Kim et al., 2017; Knockleby & Lee, 2010; Lu et al., 2009; Xu et al., 2017). We therefore performed a small interfering RNA (siRNA)-based screen targeting these MPD genes in HeLa cells to test this hypothesis. Following the transfection of cells with siRNA targeting the mRNA of each MPD gene, we immunostained the transfected cells with an antibody against Cep192, a centrosome marker, and counted the number of Cep192 foci per cell (Fig.1B,C). Among 9 MPD genes that have been implicated in DNA replication (DONSON, Orc1, Orc4, Orc6, Cdt1, Cdc6, Cdc45, GMNN, and Mcm5), we found that depletion of DONSON and Cdc6 caused a significant increase in the number of Cep192 foci (> 2) per cell (∼27%, compared to ∼1% in control cells, Fig. 1B,C). The efficacy of DONSON siRNA was confirmed by RT-qPCR analysis (Fig. S1A) since an antibody detecting endogenous DONSON protein was not available at the time the screen was carried out. Whilst it has been previously reported that Cdc6 is involved in regulating the number of centrosomes (Kim et al., 2017; Xu et al., 2017), a function of DONSON in regulating centrosome biogenesis has not been identified. We therefore focused on how DONSON regulates centrosome number.

**Fig. 1.**
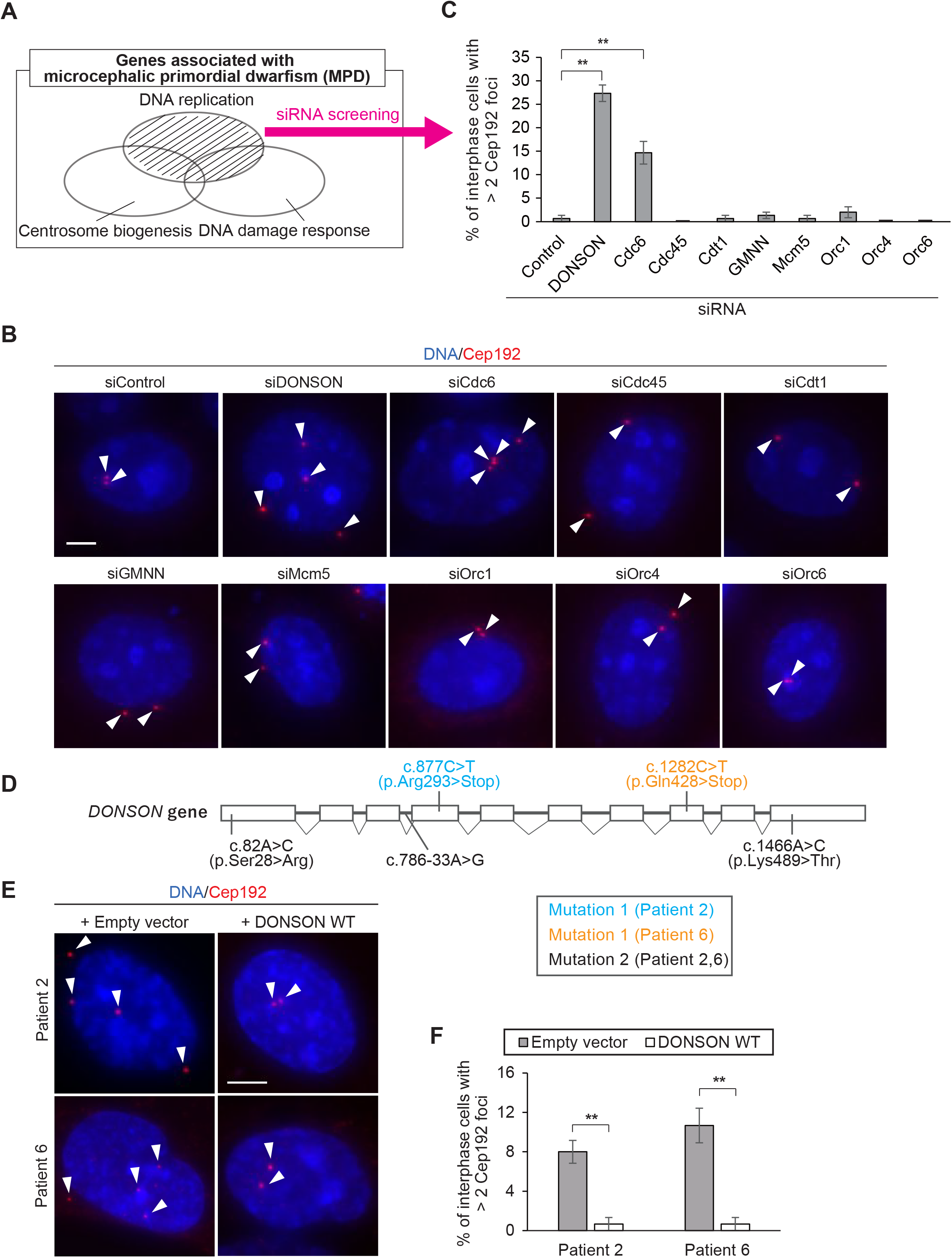
DONSON is involved in the regulation of centrosome number. **A**, List of genes involved in microcephalic primordial dwarfism (MPD) based on previous studies (Jayaraman et al., 2018). The genes involved in MPD are mostly classified as a centrosomal protein, a DNA replication component or a regulator for DNA damage response. The genes related to DNA replication pathway were submitted to siRNA screening. **B,** Aberrant number of Cep192 foci in DONSON-depleted HeLa cells. HeLa cells were treated with siControl or siRNA targeting 9 MPD genes (*DONSON, Orc1, Orc4, Orc6, Cdt1, Cdc6, Cdc45, GMNN, and Mcm5*) and immunostained with antibodies against Cep192 (red). Arrowheads indicate Cep192 foci. **C,** Histograms represent frequency of interphase cells with > 2 Cep192 foci when treated with the indicated siRNAs. Values are mean percentages ± s.d. from three independent experiments (n = 50 for each experiment). **D,** Schematic of the *DONSON* gene, indicating mutations observed in cells from MPD patient 2 (P2) and patient 6 (P6) (Reynolds et al., 2017). DONSON Patient 2 and 6 have a unique mutation in one allele (mutation 1) and both patients inherited a common haplotype containing 3 DONSON gene variants (c.82A>C (p.Ser28Arg); c.786-33A>G; c.1466A>C (p.Lys489Thr) in the other (mutation 2). The genomic structure is based on the longest ORF, containing ten coding exons (white rectangles) (NCBI NM_017613.3). **E,** Aberrant number of Cep192 foci in hTERT immortalised fibroblasts derived from patients with mutations in *DONSON*. Fibroblasts were infected with pMSCV-empty-vector or pMSCV-DONSON, and were immunostained with antibodies against Cep192 (red). Arrowheads indicate Cep192 foci. **F,** Histograms represent frequency of interphase cells with the indicated phenotypes observed in E. All scale bars, 5 μm. Tukey’s multiple comparison test was used in C, and Two-tailed, unpaired Student’s t-test was used in F to obtain P value. **, p < 0.01.

To investigate whether the existence of supernumerary centrosomes could be a cellular pathology associated with MPD in patients carrying *DONSON* mutations, we utilized hTERT immortalized, fibroblast cell lines derived from two patients with different *DONSON* mutations (Reynolds et al., 2017) (P2 and P6; Fig. 1D). Immunostaining of the cells with an antibody against Cep192 revealed that both of the patient-derived cell lines exhibited excess Cep192 foci in interphase (∼8% and ∼11%, respectively, Fig. 1E,F). This defect was complemented in both patient-derived cell lines by the expression of exogenous WT DONSON (∼1% in both cases, Fig. 1E,F), indicating that it is caused by compromising DONSON function.

We next performed live cell imaging with the HeLa cells expressing doxycycline-inducible GFP-DONSON (HeLa GFP-DONSON cells) to confirm whether DONSON localizes at the centrosome. In line with the previous immunofluorescence experiments (Giotti et al., 2018; Reynolds et al., 2017), we found that DONSON localized in the nucleus during interphase and, following nuclear envelope breakdown (NEBD), DONSON relocalized to the centrosome in mitosis (Fig. S1B,C). Next, to test whether the localization of DONSON in the nucleus or to centrosomes is necessary for its function in the regulation of centrosome number, we designed an RNAi-resistant (RNAi-R) DONSON-Full-Length (FL) construct and DONSON deletion mutant lacking the nuclear localization signal (DONSON-ΔNLS mutant) to delocalize the DONSON protein from the nucleus. Importantly, the centrosomal phenotype resulting from the siRNA-mediated depletion of endogenous DONSON was rescued by the overexpression of DONSON-FL (Fig. S1D,E), indicating that this phenotype was not due to the off-target effects. Remarkably, however, that was not the case with the DONSON-ΔNLS mutant (Fig. S1D,E). Therefore, these data suggest that DONSON is involved in maintaining proper centrosome number and that displacement of DONSON from the nucleus compromises this function.

### Depletion of DONSON causes precocious centriole disengagement in interphase

To gain insight into how depletion of DONSON leads to aberrant number of centrosomes, we monitored the number of centrioles by immunostaining DONSON-depleted cells with antibodies against centrin-2, a centriole marker. As expected, cells transfected with control siRNA mostly harbored two pairs of centrin foci, indicating two mother-daughter centriole pairs in S and G2 phase (Fig. 2A,B). However, in contrast, we found that DONSON-depleted cells exhibited four separated centrin foci in interphase (26%, Fig. 2A,B), and that these centrin foci were predominantly marked with Cep192. The same phenotype was also observed when using siRNA targeting a different sequence of the DONSON ORF (Fig. S2). This result raised the possibility that precocious centriole disengagement was occurring in interphase cells, upon depletion of DONSON.

**Fig. 2.**
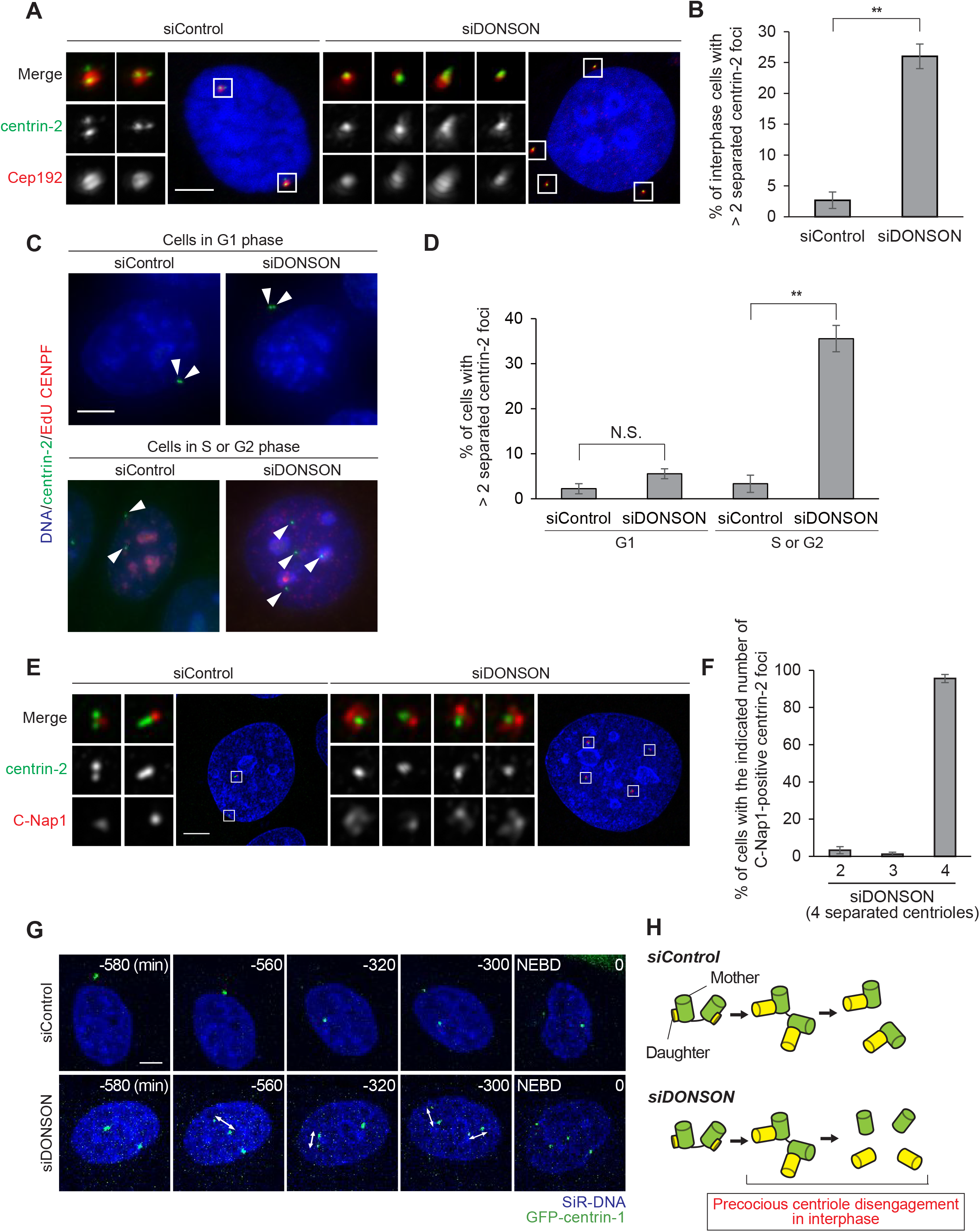
Depletion of DONSON causes centriole disengagement in interphase. **A**, Depletion of DONSON induced four separated centrioles in HeLa cells. HeLa cells were treated with siControl or siDONSON and immunostained with antibodies against centrin-2 (green) and Cep192 (red). **B,** Histograms represent frequency of interphase cells with the indicated phenotypes observed in A. Values are mean percentages ± s.d. from three independent experiments (n = 50 for each experiment). **C,** Depletion of DONSON did not affect the number of centrin foci in G1 phase. HeLa cells were treated with siControl or siDONSON for 48 h, and 10 μM EdU for 30 min before fixation, followed by immunostaining with antibodies against centrin-2 (green) and CENP-F (red). S phase cells were monitored by EdU staining (red). CENPF and EdU double-negative cells were classified as G1 phase cells, and others as S or G2 phase cells. **D,** Histograms represent frequency of interphase cells with the indicated phenotype observed in C. Values are mean percentages ± s.d. from three independent experiments (n = 30 for each experiment). **E,** Recruitment of C-Nap1 to separated centrioles in DONSON-depleted HeLa cells. HeLa cells were treated with siControl or siDONSON and immunostained with antibodies against centrin-2 (green) and C-Nap1 (red). **F,** Histograms represent frequency of cells with the indicated number of C-Nap1 positive centrioles among cells with four separated centrioles observed in E. Values are mean percentages ± s.d. from three independent experiments (n = 30 for each experiment). **G,** Time-lapse observation of precocious centriole disengagement in interphase in DONSON-depleted cells. HeLa GFP-centrin-1 cells were visualized every 20 min for 48 h after 24 h treatment of siControl or siDONSON and 3 h treatment of 100 nM SiR-DNA. Left– right arrows indicate precociously disengaged centrioles. **H,** Schematic model of the phenotypes of DONSON depletion. Depletion of DONSON causes precocious centriole disengagement in interphase. All scale bars, 5 μm. Two-tailed, unpaired Student’s t-test was used in B and D to obtain P value. **, p < 0.01; N.S., not significantly different (p > 0.05).

Generally, the appearance of four separated centrioles in interphase can arise from a cytokinesis failure or precocious centriole disengagement. Therefore, we first examined the former possibility in DONSON-depleted cells. Since cells that fail to undergo cytokinesis in the previous cell cycle generally exhibit four separated centrioles in G1 phase, we examined the number of separated centrioles in cells lacking DONSON in different phases of the cell cycle. Notably, we observed that most DONSON-depleted cells exhibiting 4 separated centrioles were in S/G2 phase rather than G1 phase of the cell cycle (Fig. 2C,D). These data suggest that the supernumerary centrosome phenotype seen in DONSON-depleted cells was not due to a cytokinesis failure. Consistently, we found that C-Nap1, a protein which localizes at the proximal end of mother and disengaged daughter centrioles (Mayor, Stierhof, Tanaka, Fry, & Nigg, 2000; Yang, Adamian, & Li, 2006) was recruited to all of the separated centrioles during interphase in DONSON-depleted cells (∼96%, Fig. 2E,F). Moreover, live cell imaging of HeLa cells stably expressing GFP-centrin-1 revealed that, when DONSON is depleted, a pair of centrioles disengaged in interphase and then dynamically moved apart from its mother centriole (Fig. 2G). Collectively, these data strongly suggest that depletion of DONSON caused precocious centriole disengagement in interphase (Fig. 2H).

### Depletion of DONSON leads to centriole-to-centrosome conversion of disengaged daughter centrioles and centriole reduplication

We previously demonstrated that precociously-disengaged centrioles aberrantly acquire PCM components and ectopic MTOC activity in early mitosis (Watanabe, Takao, Ito, Takahashi, & Kitagawa, 2019). Therefore, we investigated whether disengaged daughter centrioles in DONSON-depleted cells are converted to centrosomes in interphase. Immunostaining with antibodies against PCNT and γ-Tubulin (major PCM components) revealed that almost all of the disengaged centrioles in DONSON-depleted cells recruited PCM to their periphery (∼91%, Fig. 3A,B,S3A). We next performed microtubule regrowth assay to confirm whether the disengaged centrioles acquire ectopic MTOC activity in DONSON-depleted cells. For this analysis, we first depolymerized the microtubule network using a combined treatment of cold shock and nocodazole. Subsequently, we returned the cells to permissive conditions and allowed centrosome-organized microtubules to re-grow (Fig. 3C). This assay demonstrated that in DONSON-depleted cells, precociously disengaged centrioles obtained ectopic MTOC activity in interphase (Fig. 3C).

**Fig. 3.**
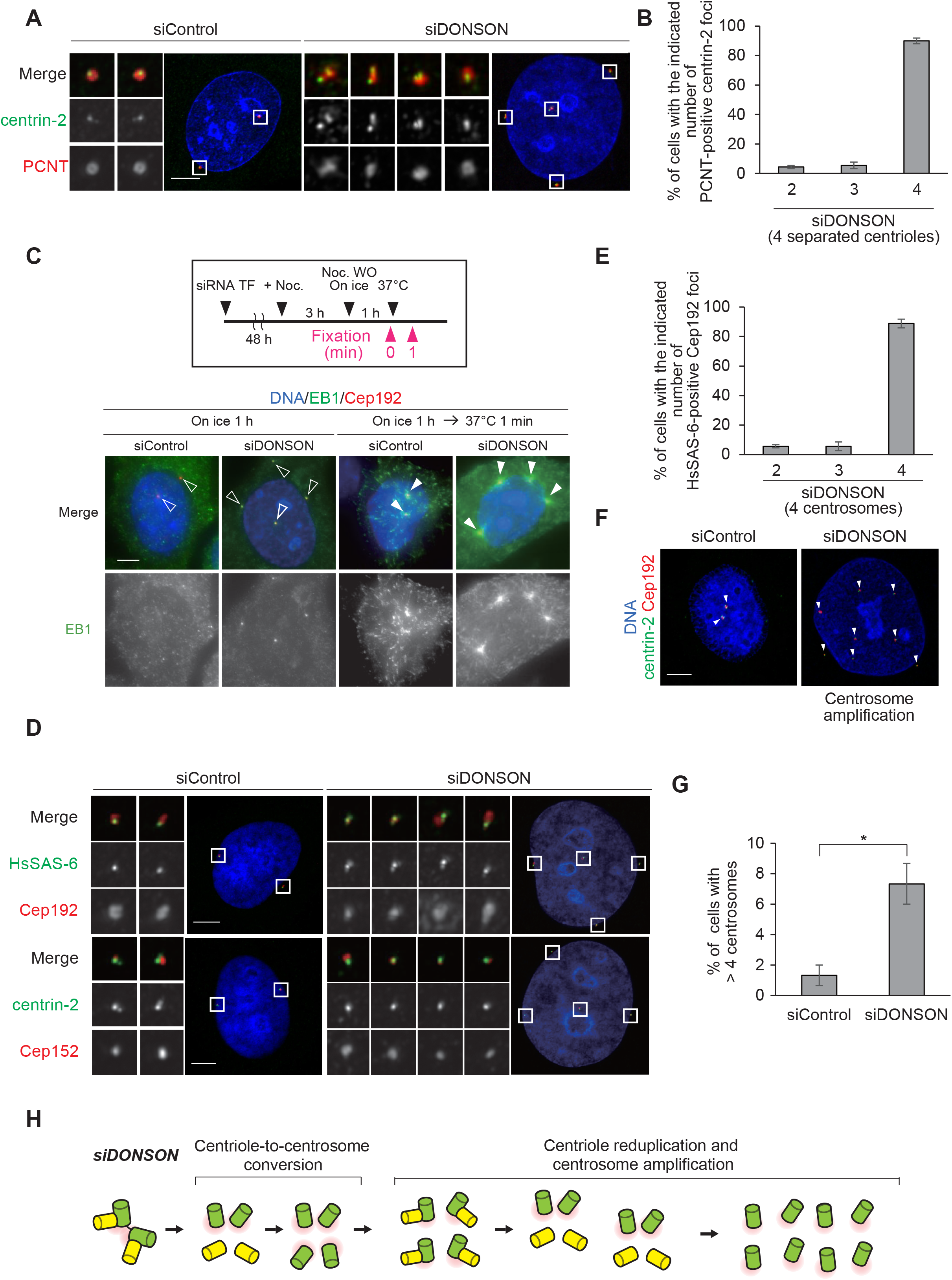
Centriole-to-centrosome conversion and centriole reduplication in DONSON-depleted cells. **A**, Aberrant acquisition of PCM components by precociously disengaged daughter centrioles in DONSON-depleted cells. HeLa cells were immunostained with antibodies against centrin-2 (green) and PCNT (red). **B,** Histograms represent frequency of DONSON-depleted cells with the indicated number of PCNT foci among cells with four separated centrioles in interphase. Values are mean percentages ± s.d. from three independent experiments (n = 30 for each experiment). **C,** Microtubule regrowth assay after 10 μM Nocodazole treatment for 3 h and subsequent ice-cold treatment for 1 h in DONSON-depleted cells. HeLa cells were immunostained with antibodies against EB1 (green) and Cep192 (red). White arrowheads indicate centrosomes with MTOC activity. Arrowheads with white outline indicate centrosomes without microtubule nucleation. **D,** Centriole reduplication after depletion of DONSON in HeLa cells. HeLa cells were treated with siControl or siDONSON for 48 h and immunostained with the indicated antibodies. **E,** Histograms represent frequency of interphase cells with the indicated number of HsSAS-6 foci among cells with four separated centrioles observed in D. Values are mean percentages ± s.d. from three independent experiments (n = 50 for each experiment). **F,** HeLa cells were treated with siControl or siDONSON for 48 h and immunostained with antibodies against centrin-2 (green) and Cep192 (red). Arrowheads indicate Cep192 foci (centrosomes). **G,** Histograms represent frequency of interphase cells with the indicated phenotypes observed in F. Values are mean percentages ± s.d. from three independent experiments (n = 50 for each experiment). Two-tailed, unpaired Student’s t-test was used to obtain P value. *, p < 0.05. **H,** Schematic model of the phenotype of DONSON depletion observed in A, D and F. Depletion of DONSON led to centriole disengagement, centriole-to-centrosome conversion, acquisition of ectopic MTOC activity and centriole amplification in interphase. All scale bars, 5 μm.

Since disengaged daughter centrioles aberrantly recruited PCM components in interphase in DONSON-depleted cells, we investigated whether the disengaged centrioles are converted to centrosomes harboring the ability to duplicate (Fu et al., 2016; Izquierdo, Wang, Uryu, & Tsou, 2014; Wang, Soni, Uryu, & Tsou, 2011). Considering the previous report showing that centriole engagement blocks reduplication of centrioles (Loncarek et al., 2008; Tsou et al., 2009; Tsou & Stearns, 2006), we hypothesized that precocious centriole disengagement would lead to reduplication of a new centriole (Lončarek, Hergert, & Khodjakov, 2010). Since Cep152 forms scaffolds for centriole duplication by recruiting Plk4 (Cizmecioglu et al., 2010; Hatch, Kulukian, Holland, Cleveland, & Stearns, 2010), we immunostained DONSON-depleted cells with an antibody against Cep152. Consistent with our hypothesis, we found that Cep152 was recruited to the disengaged daughter centrioles in DONSON-depleted cells (Fig. 3D). We next asked whether precociously-disengaged mother and daughter centrioles acquire the ability to duplicate a new centriole. To do this, we performed immunostaining with an antibody against HsSAS-6, a procentriole marker, and found that HsSAS-6 localized at almost all of the disengaged centrioles in DONSON-depleted cells (∼91%, Fig. 3D,E). Consistently, we often found aberrant number of centrosomes, marked by Cep192, harboring a pair of centrioles foci in DONSON-depleted cells (Fig. S3B). Moreover, a previous study suggested that centriole disengagement and subsequent centriole reduplication in interphase can be a repetitive process, which leads to accelerated centrosome amplification (Lončarek et al., 2010). We therefore investigated whether centrioles which were newly reduplicated after centriole disengagement in DONSON-depleted cells also lead to centrosome amplification. Compared with control cells, we more frequently observed cells with more than four centrosomes following DONSON-depletion, indicating that precocious centriole disengagement and associated centriole reduplication promotes centrosome amplification in DONSON-depleted cells (∼7%, compared with ∼1.5% in control cells, Fig. 3F,G).

Collectively, these data suggest that in interphase, DONSON prevents centriole reduplication caused by centrosome conversion of precociously disengaged centrioles (Fig. 3H).

### Amplified centrosomes are clustered during interphase through the p53 pathway in DONSON-depleted cells

To investigate whether the precocious centriole disengagement phenotype observed following DONSON depletion was not specific to HeLa cells, we transfected different human cell lines such as U2OS, PANC-1 and RPE-1, with DONSON siRNA and examined centriole behavior using immunofluorescence. Although depletion of DONSON induced precocious centriole disengagement in all cell lines that we tested, interestingly, we found that the distance between disengaged centrioles was different in interphase, depending on the cell line (Fig. 4A,B,S4A,B). Therefore, we quantified the distance between a pair of disengaged centrioles in each DONSON-depleted cell line according to whether the distance was more or less than 2.5 μm for simplicity. In HeLa and PANC-1 cells, the distance between a pair of disengaged centrioles was predominantly more than 2.5 μm (∼75% and ∼80%, respectively), and they were often dispersed in the cytoplasm. In contrast, in U2OS and RPE-1 cells, the distance was largely less than 2.5 μm (∼73% and ∼83%, respectively), and a pair of mother and daughter centrioles were usually still in close proximity even after they were disengaged. Interestingly, we noticed that these two cell groups can be distinguished by the functional status of the p53 pathway. Consistent with p53 playing a role in determining the distance between disengaged mother and daughter centrioles, it has been previously shown that centrosome clustering is mediated by the p53 pathway in tetraploid mouse embryonic fibroblasts (Yi et al., 2011). We therefore hypothesized that the difference in the distance of disengaged centrioles may be due to whether or not the cell line retains an intact p53 pathway. To address this, we investigated whether depletion of p53 impaired the clustering of supernumerary centrosomes during interphase in DONSON-depleted U2OS cells. Consistent with our hypothesis, co-depletion of DONSON and p53, using two different siRNA sequences inhibited the clustering of disengaged centrioles in cells lacking DONSON (Fig. 4C,D,S4C). It has been shown that p53-dependent centrosome clustering is inhibited by activation of actin severing protein, cofilin. To further test that the p53 pathway-dependent centrosome clustering machinery acts to suppress the separation of supernumerary centrosomes in DONSON-depleted cells, we disturbed centrosome clustering by cofilin-mediated disruption of the cortical actin cytoskeleton, using Crenolanib. As expected, we found that treatment of Crenolanib induced separation of supernumerary centrosomes during interphase in DONSON-depleted U2OS cells (∼74%, compared to ∼27% in control (DMSO), Fig. 4E,F). Taken together, supernumerary centrosomes from precocious centriole disengagement in DONSON-depleted cells are clustered in interphase by p53-dependent centrosome clustering machinery (Fig. 4G).

**Fig. 4.**
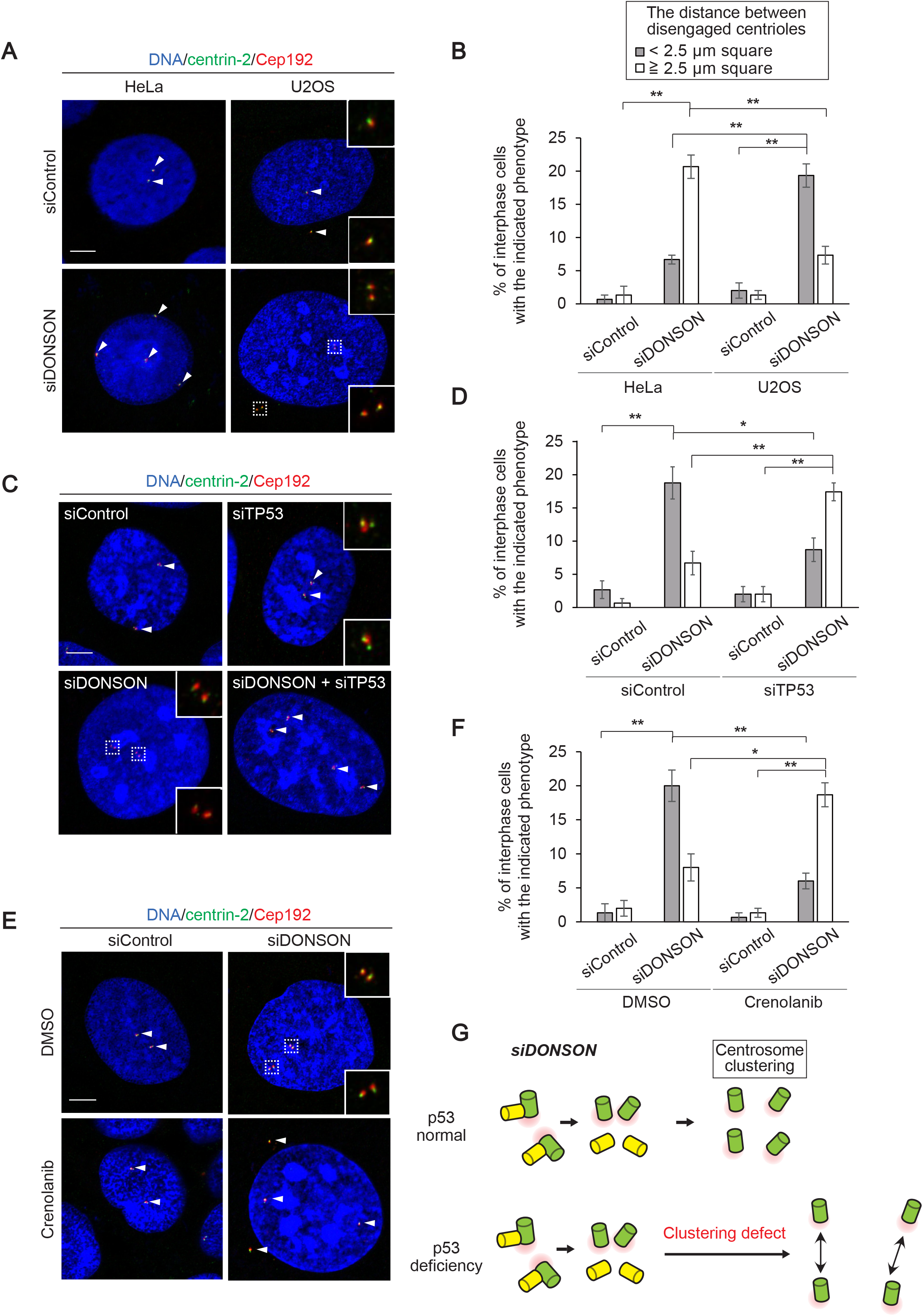
p53-dependent centrosome clustering in DONSON-depleted cells. **A**, Centriole disengagement caused by DONSON depletion in HeLa and U2OS cells. The cells were immunostained with antibodies against centrin-2 (green) and Cep192 (red). Arrowheads indicate Cep192 foci (centrosomes). **B,** Histograms represent frequency of interphase cells with the indicated phenotypes observed in A. Values are mean percentages ± s.d. from three independent experiments (n = 50 for each experiment). **C,** p53-dependent centrosome clustering in DONSON-depleted U2OS cells. U2OS cells were treated with the indicated siRNAs for 48 h and immunostained with antibodies against centrin-2 (green) and Cep192 (red). Arrowheads indicate Cep192 foci (centrosomes). **D,** Histograms represent frequency of interphase cells with the indicated phenotypes observed in C. Values are mean percentages ± s.d. from three independent experiments (n = 50 for each experiment). **E,** Disruption of cortical actin cytoskeleton reduced centrosome clustering in U2OS cells. U2OS cells were treated with siControl or siDONSON for 48 h and DMSO or Crenolanib for 48 h. Arrowheads indicate Cep192 foci (centrosomes). **F,** Histograms represent frequency of interphase cells with the indicated phenotypes observed in E. Values are mean percentages ± s.d. from three independent experiments (n = 50 for each experiment). **G,** Schematic model of p53-dependent centrosome clustering after ectopic centrosome conversion in DONSON-depleted cells. All scale bars, 5 μm. All the squares with white dotted lines are 2.5 μm square. Tukey’s multiple comparison test was used in B, D, and F to obtain P value. *, p < 0.05; **, p < 0.01.

### Defects in mitotic spindle formation and chromosome segregation in DONSON-depleted cells

Since precocious centriole disengagement causes chromosome segregation errors in mitosis (Watanabe et al., 2019), we next focused on the impact of DONSON depletion on mitotic cell division. To observe how disengaged centrioles affect mitotic spindle formation and chromosome segregation, we performed live-cell imaging on HeLa cells stably expressing GFP-centrin-1 following transfection with DONSON siRNA. Importantly, we found that DONSON-depleted cells frequently exhibited mitotic defects, such as chromosome misalignment with pseudo-bipolar spindle formation and multi-polar spindle formation (∼19% and ∼30%, respectively, Fig. 5A,B), which were frequently associated with lagging chromosomes and/or multi-nucleated cells. Interestingly, we noted that these abnormal spindle formations arose from disengaged centrioles which had acquired ectopic MTOC activity in interphase (Fig. 3C). Similar defects were also confirmed in DONSON-depleted fixed cells (Fig. S5A,B). In addition, we noticed a higher frequency of multi-polar spindle formation in HeLa cells than U2OS cells (Fig. 5C,D,E,F). Given that supernumerary centrosomes derived from precocious centriole disengagement were clustered in interphase through the p53 pathway in U2OS cells (Fig. 4A,B), we investigated whether p53 deficiency affects the frequency of multi-polar spindle formation in DONSON-depleted U2OS cells. To address this, we co-depleted DONSON and p53 in U2OS cells, and monitored spindle formation during mitosis using fluorescence microscopy. Consistently, we observed that the combined depletion of DONSON and p53 increased the frequency of multi-polar spindle formation in U2OS cells (∼21%, compared to ∼9% upon DONSON depletion, Fig. 5E,F). In keeping with this, multi-polar cell division was also induced by direct inhibition of centrosome clustering using crenolanib (Fig. S5C,D). Together, these results suggest that the p53 pathway suppresses multi-polar spindle formation in DONSON-depleted cells by clustering extra centrosomes.

**Fig. 5.**
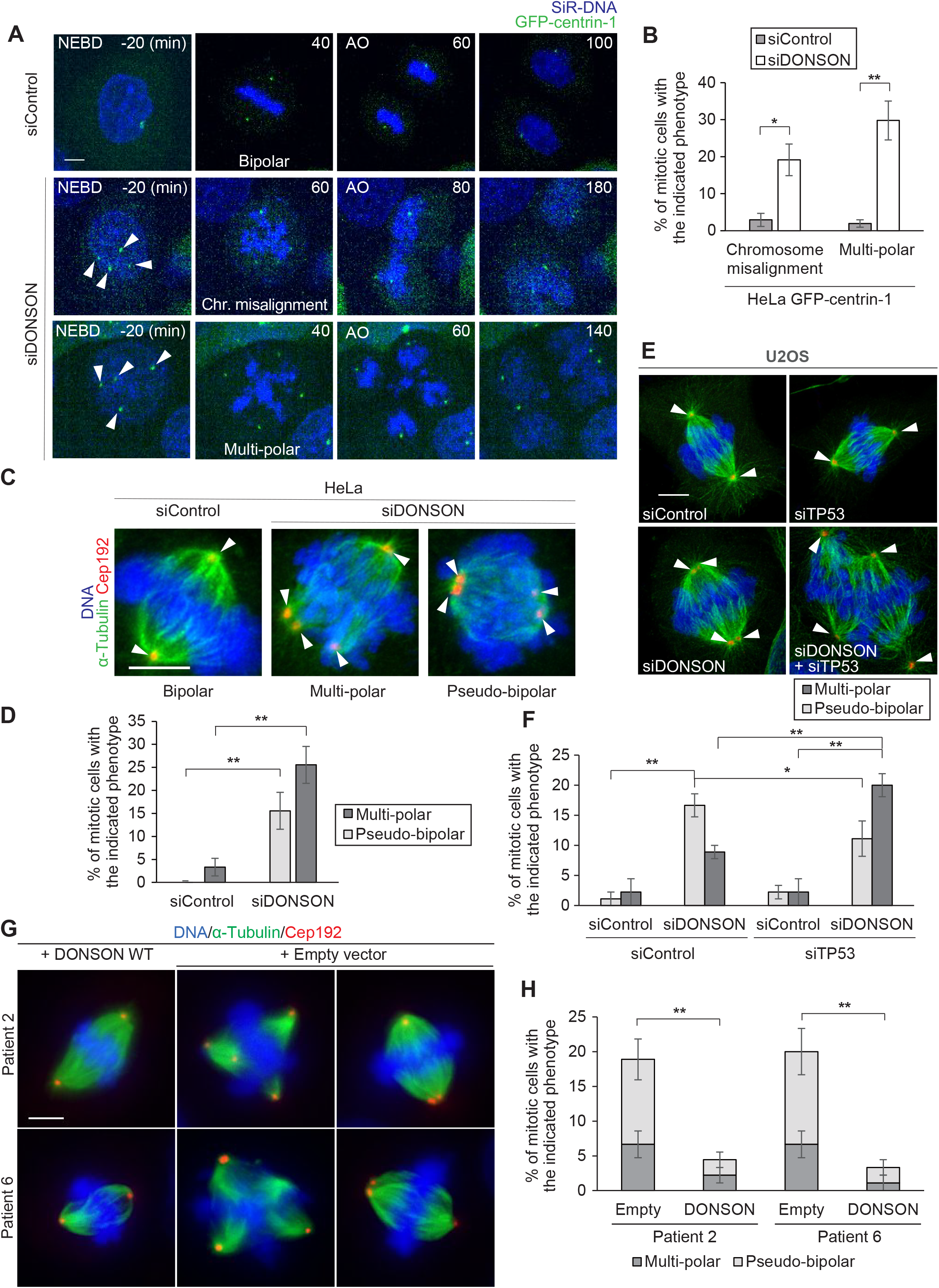
Defects in mitotic spindle formation and chromosome segregation in DONSON-depleted cells. **A**, Mitotic defects observed in DONSON-depleted cells. HeLa GFP-centrin-1 cells were treated with siControl or siDONSON in the presence of SiR-DNA (100 nM). Arrowheads indicate precociously disengaged centrioles. **B,** Histograms represent frequency of the mitotic cells with the indicated phenotypes observed in A (siControl n = 106, siDONSON n = 94 from three independent experiments). **C,** Abnormal spindle formations in DONSON-depleted HeLa cells. HeLa cells were treated with siControl or siDONSON and immunostained with antibodies against α-Tubulin (green) and Cep192 (red). Arrowheads indicate Cep192 foci (centrosomes). **D,** Histograms represent frequency of mitotic cells with the indicated phenotypes observed in C. Values are mean percentages ± s.d. from three independent experiments (n = 30 for each experiment). **E,** Abnormal spindle formations in DONSON-depleted U2OS cells. U2OS cells were treated with siControl, siDONSON, siTP53 or siDONSON and siTP53 and were immunostained with antibodies against α-Tubulin (green) and Cep192 (red). Arrowheads indicate Cep192 foci (centrosomes). **F,** Histograms represent frequency of mitotic cells with the indicated phenotypes observed in E. Values are mean percentages ± s.d. from three independent experiments (n = 30 for each experiment). **G,** Abnormal spindle formations in fibroblasts derived from patients with mutations in *DONSON*. The cells were immunostained with antibodies against α-Tubulin (green) and Cep192 (red). **H,** Histograms represent frequency of mitotic cells with the indicated phenotypes observed in G. Values are mean percentages ± s.d. from three independent experiments (n = 30 for each experiment). All scale bars, 5 μm. Two-tailed, unpaired Student’s t-test was used in B, D, and H, and Tukey’s multiple comparison test was used in F to obtain P value. *, p < 0.05; **, p < 0.01.

Lastly, we characterized mitotic spindle formation in patient-derived cell lines with *DONSON* mutations. As expected, these cells showed pseudo-bipolar or multi-polar spindle formation (∼15% and ∼7%, respectively, in Patient 2, ∼18% and ∼7%, respectively, in Patient 6, Fig. 5G,H). Again, this phenotype was complemented by the re-expression of DONSON WT (Fig. 5G,H), suggesting that these defects were directly caused by a functional deficiency of DONSON. Overall, these findings strongly suggest DONSON is involved in maintaining proper centrosome number, and thereby proper spindle formation and chromosomal segregation.

## Discussion

In this study, we addressed the potential function of DONSON in centrosome biogenesis (Fig. 6). We revealed that depletion of DONSON causes precocious centriole disengagement and subsequent centriole reduplication in interphase. These defects in the centriole duplication cycle leads to a failure of the mitotic process, such as multi-polar spindle formation and chromosome segregation errors. Importantly, cells derived from patients with *DONSON* mutations also exhibited precocious centriole disengagement and accompanying mitotic defects, suggesting this cellular phenotype may contribute to the neurodevelopmental dysfunction exhibited by the affected patients.

**Fig. 6.**
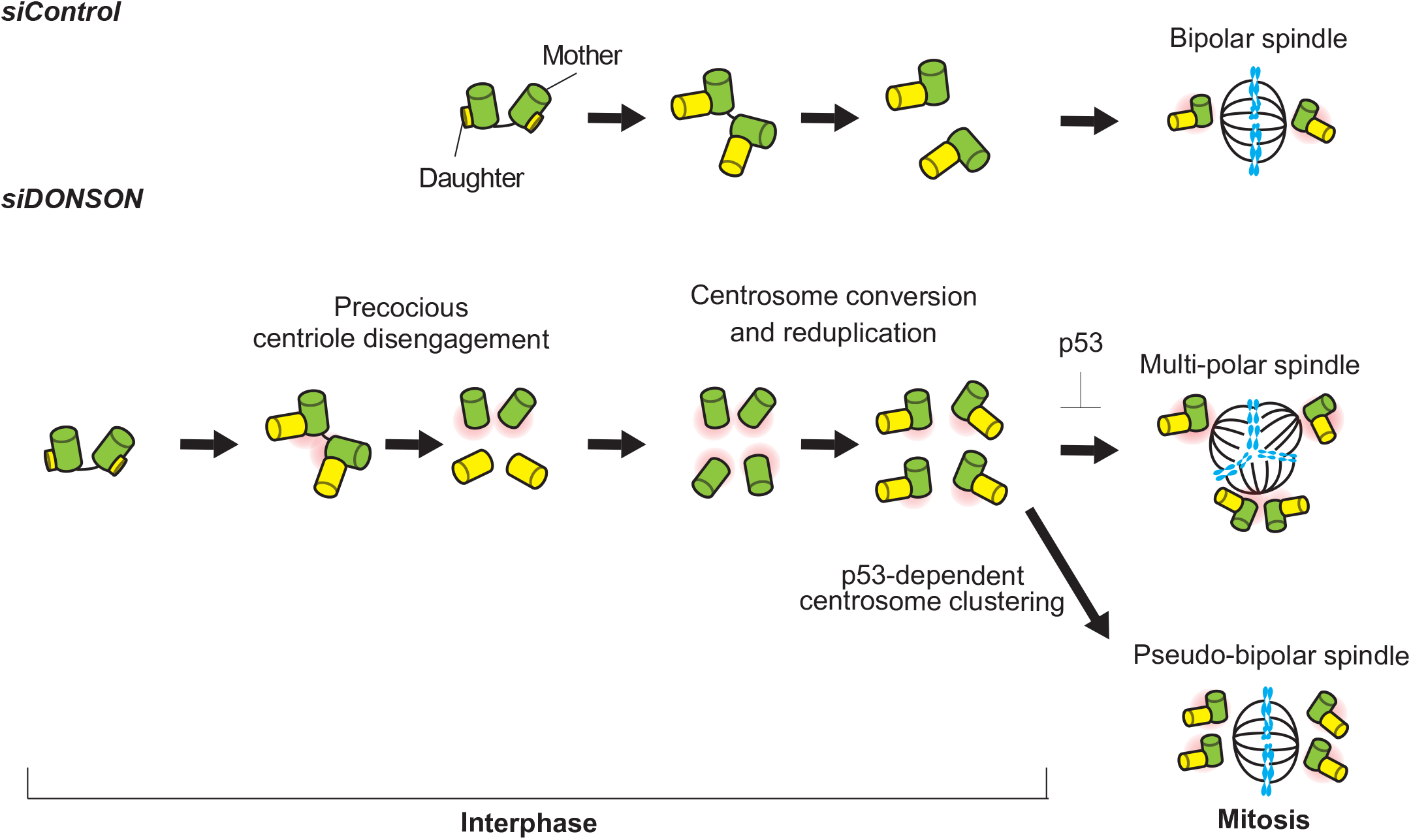
A schematic model of how depletion of DONSON affects centriole biogenesis. Depletion of DONSON causes precocious centriole disengagement in interphase. Precociously disengaged daughter centrioles aberrantly acquire PCM components and convert to centrosomes without passing through mitosis. Since converted centrosomes ectopically acquire MTOC activity, DONSON-depleted cells exhibit abnormal spindle formation and chromosome segregation errors in mitosis. In particular, extra centrosomes arose by DONSON depletion are clustered by p53-dependent pathway, and mutation or deficiency of p53 induces multi-polar spindle formation and aneuploidy more frequently in DONSON-depleted cells.

Previously, two studies identified *DONSON* as a novel causative gene of MPD (Evrony et al., 2017; Reynolds et al., 2017). Whilst, DONSON has been reported to be a component of the DNA replication fork (Reynolds et al., 2017), it is known that other replisome components linked to MPD, such as Orc1, Cdc6, and GMNN, also localize to centrosomes (Hemerly et al., 2009; Kim et al., 2017; Knockleby & Lee, 2010; Lu et al., 2009; Xu et al., 2017), suggesting that specific subunits of the replication machinery may have dual roles, one during S phase and one during mitosis. Consistent with this premise, we confirmed that DONSON localized in the nucleus during interphase and centrosomes during mitosis (Fig. S1B,C). Interestingly however, we demonstrated that the precocious centriole disengagement and reduplication observed in DONSON-depleted cells were due to a loss of functional DONSON from the nucleus rather than the centrosomes (Fig. S1D,E). This suggests that there is a potential link between DNA replication and the regulation of centriole engagement and disengagement although the underlying mechanism remains to be determined.

A previous study demonstrated that, upon depletion of Cep57, centrioles are precociously disengaged in early mitosis, and that these aberrantly disengaged daughter centrioles recruit PCM components and acquire ectopic MTOC activity, which ultimately causes chromosome segregation errors and aneuploidy during the transit through mitosis (Watanabe et al., 2019). We also observed similar phenotypes in DONSON-depleted cells (Fig. 5A,B,S5A,B). However, in addition to the acquisition of ectopic MTOCs, centriole disengagement that occurs in interphase has more harmful effects. Since the disengaged centrioles caused by DONSON depletion can reduplicate within the same interphase (Fig. 3D,E), this phenotype is also accompanied with numerical centrosome aberrations. These extra centrosomes could have various effects on human cells other than mitotic defects and aneuploidy, which are related to carcinogenesis. For example, extra centrosomes can compromise cell signaling by affecting cilia formation, and cell polarity by increasing microtubule nucleation (Godinho & Pellman, 2014). Thus, precocious centriole disengagement in interphase is likely to be more detrimental for cells in all phases of the cell cycle than just during mitosis. Therefore, further investigation to understand how depletion of DONSON leads to precocious centriole disengagement will be an important issue for future study.

Although *DONSON* has been identified as a causative gene of MPD, how mutations in *DONSON* lead to the phenotypes of MPD patients still remains elusive. A previous study reported spontaneous defects in replication fork stability and increased levels of DNA damage in patient cells with *DONSON* mutations, which were suggested as one possible cause of MPD (Reynolds et al., 2017). Meanwhile, we also found precocious centriole disengagement in interphase and accompanying mitotic defects in patient cells with *DONSON* mutations (Fig.1E,F,5G,H). It has been suggested that precocious centriole disengagement could represent a potential underlying cause of microcephaly. Previous studies indicated that depletion of Cep57 or PCNT causes precocious centriole disengagement in early mitosis and accompanying chromosome segregation errors (Lee & Rhee, 2012; Matsuo et al., 2012; Watanabe et al., 2019). Indeed, MVA patient cells with *CEP57* mutations exhibited precocious centriole disengagement and chromosome segregation errors (Watanabe et al., 2019). Notably, mutations in both *CEP57* and *PCNT* have been associated with the development of microcephaly (Rauch et al., 2008; Snape et al., 2011). Thus, whilst precocious centriole disengagement may represent a contributory factor for neurodevelopmental disease, it is not clear how differences in the timing of centriole disengagement i.e. interphase versus mitosis, affects disease severity or the presence/absence of additional clinical phenotypes e.g. hypoplastic limbs in patients with DONSON associated microcephaly-micromelia syndrome.

It is assumed that the activity of Plk1, APC/C (anaphase-promoting complex/cyclosome) and separase, which are known to be normally involved in centriole disengagement during mitosis, is relatively low during interphase. Therefore, it will be important to clarify, in future study, how compromised function of DONSON leads to precocious centriole disengagement even in interphase.

## Contribution

K.M. and D.K. designed the study; K.M., K.W., T.C., S.H. and D.K. designed experiments; K.M. and K.K.I. performed experiments; K.M., K.K.I., K.W., S.H., and D.K. analyzed data; G.S.S generated the DONSON patient cell lines; K.M. and D.K. wrote the manuscript, which was commented on by all authors.

## Acknowledgements

We gratefully acknowledge the members of the Kitagawa laboratory for technical support and discussion. This work was supported by a Grant-in-Aid for Scientific Research (S, 19H05651) and Research Activity start-up from the Ministry of Education, Science, Sports and Culture of Japan, by the Takeda Science Foundation, by the Japan Science Society (The Sasagawa Scientific Research Grant), by the Daiichi Sankyo Foundation of Life Science. G.S.S. is supported by a Cancer Research UK (CR-UK) programme grant (C17183/A23303).

---

**Fig. S1.**
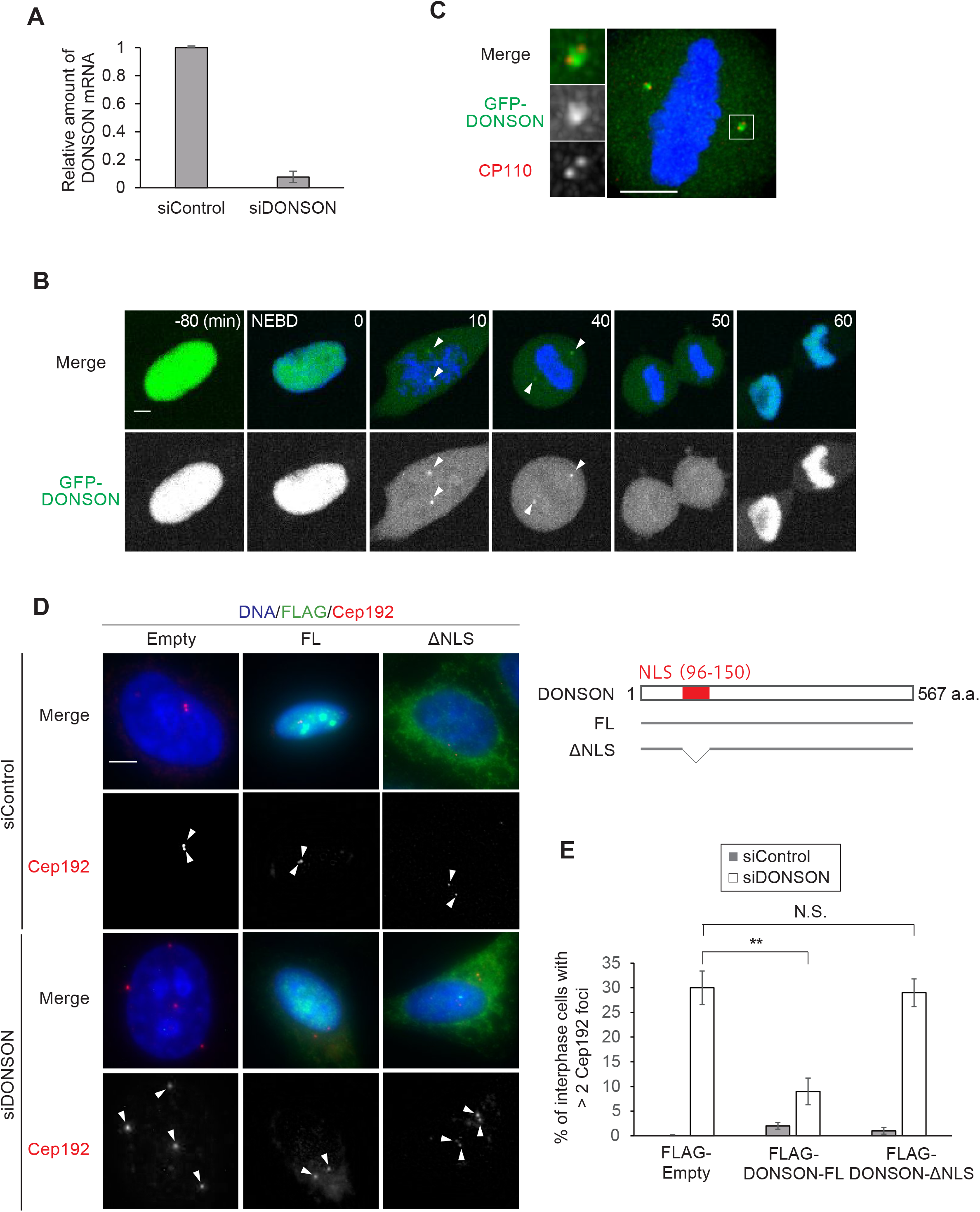
**A**, Quantitative PCR (qPCR) analysis of DONSON mRNA amount in HeLa cells. Total RNA was isolated from HeLa cells after 48 h treatment of the indicated siRNAs, and was subjected to reverse transcription. The resulting cDNA was subjected to real-time qPCR analysis. **B,** Time-lapse observation of DONSON localization at different cell cycle stages. HeLa GFP-DONSON cells were visualized every 10 min for 24 h after 24 h treatment of 1 μg/mL Doxycycline and 3 h treatment of 100 nM SiR-DNA. DNA (blue) and GFP-DONSON (green). Arrowheads indicate GFP-DONSON foci. **C,** The centrosomal localization of DONSON in mitosis. HeLa GFP-DONSON cells were fixed after 24 h treatment of 1 μg/mL Doxycycline and were immunostained with antibodies against GFP (green) and CP110 (red). **D,** Phenotype rescue experiments by expression of RNAi-resistant (RNAi-R) form of full-length (FL) or nuclear localization signal deleted (ΔNLS) DONSON. HeLa cells were treated with siControl or siDONSON, followed by transfection with FLAG empty (control), RNAi-resistant DONSON-FL, or DONSON-ΔNLS. The cells were immunostained with antibodies against FLAG (green) and Cep192 (red). Arrowheads indicate Cep192 foci. **E,** Histograms represent frequency of interphase cells with the indicated phenotypes observed in D. Values are mean percentages ± s.d. from three independent experiments (n = 30 for each experiment). Tukey’s multiple comparisons test was used in E to obtain P value. **, p < 0.01; N.S., not significantly different (p > 0.05). All scale bars, 5 μm.

**Fig. S2.**
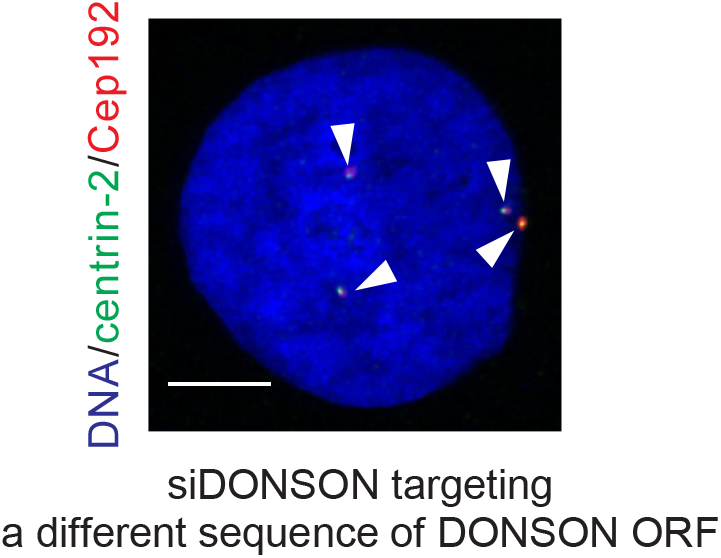
The phenotype induced by DONSON depletion was confirmed by using another siRNAHeLa cells were treated with siRNAs targeting against different sequences of DONSON ORF and were immunostained with antibodies against centrin-2 (green) and Cep192 (red). Arrowheads indicate Cep192 foci. Scale bar, 5 μm.

**Fig. S3.**
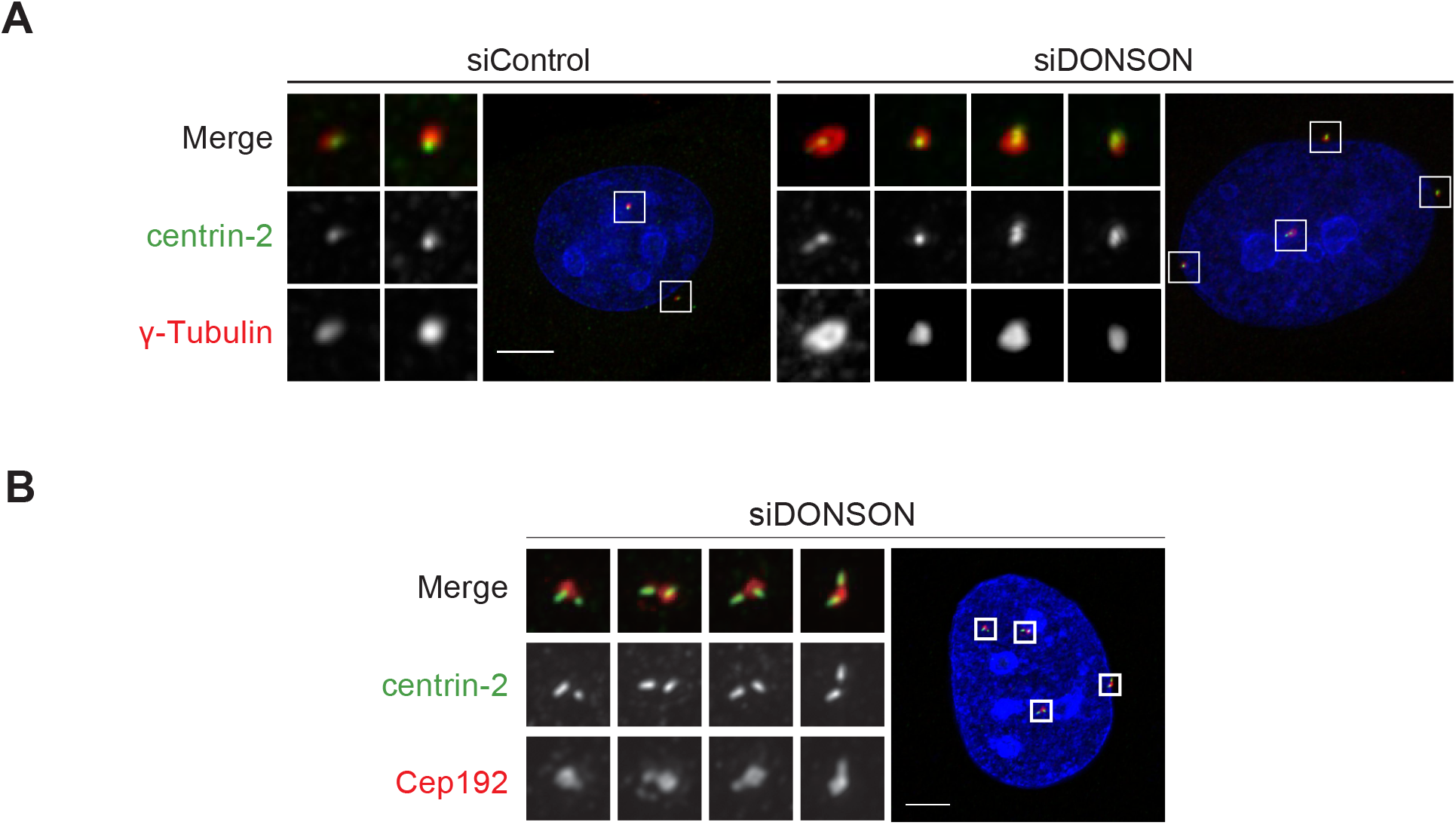
**A**, Ectopic acquisition of PCM component after depletion of DONSON in HeLa cells. HeLa cells were treated with siControl or siDONSON for 48 h and immunostained with antibodies against centrin-2 (green) and γ-Tubulin (red). **B,** Cells with aberrant number of diplosomes after DONSON depletion. HeLa cells were treated with siRNA for 48 h and were immunostained with antibodies against centrin-2 (green) and Cep192 (red). All scale bars, 5 μm.

**Fig. S4.**
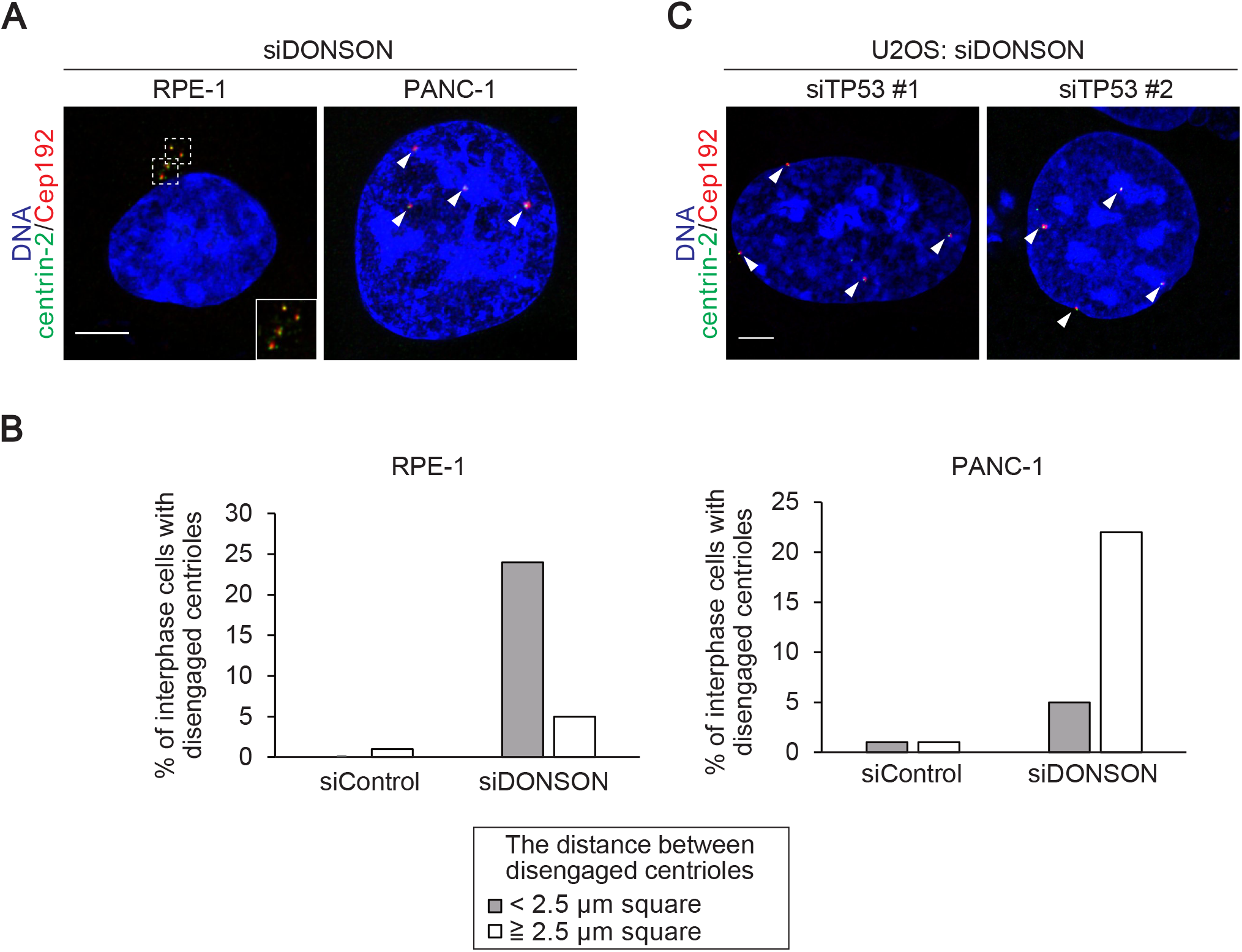
**A**, The distance between disengaged centrioles was different in interphase, depending on the cell lines. RPE-1 and PANC-1 cells were treated with siDONSON and immunostained with antibodies against centrin-2 (green) and Cep192 (red). Arrowheads indicate precociously disengaged centrioles. **B,** Histograms represent frequency of interphase cells with the indicated phenotype observed in A. (two independent experiments, N=50 for each experiment). **C,** The phenotype induced by p53 depletion was confirmed by using another siRNA. U2OS cells were treated with DONSON siRNA and siRNAs targeting against different sequences of TP53 ORF and were immunostained with antibodies against centrin-2 (green) and Cep192 (red). Arrowheads indicate precociously disengaged centrioles. All scale bars, 5 μm.

**Fig. S5.**
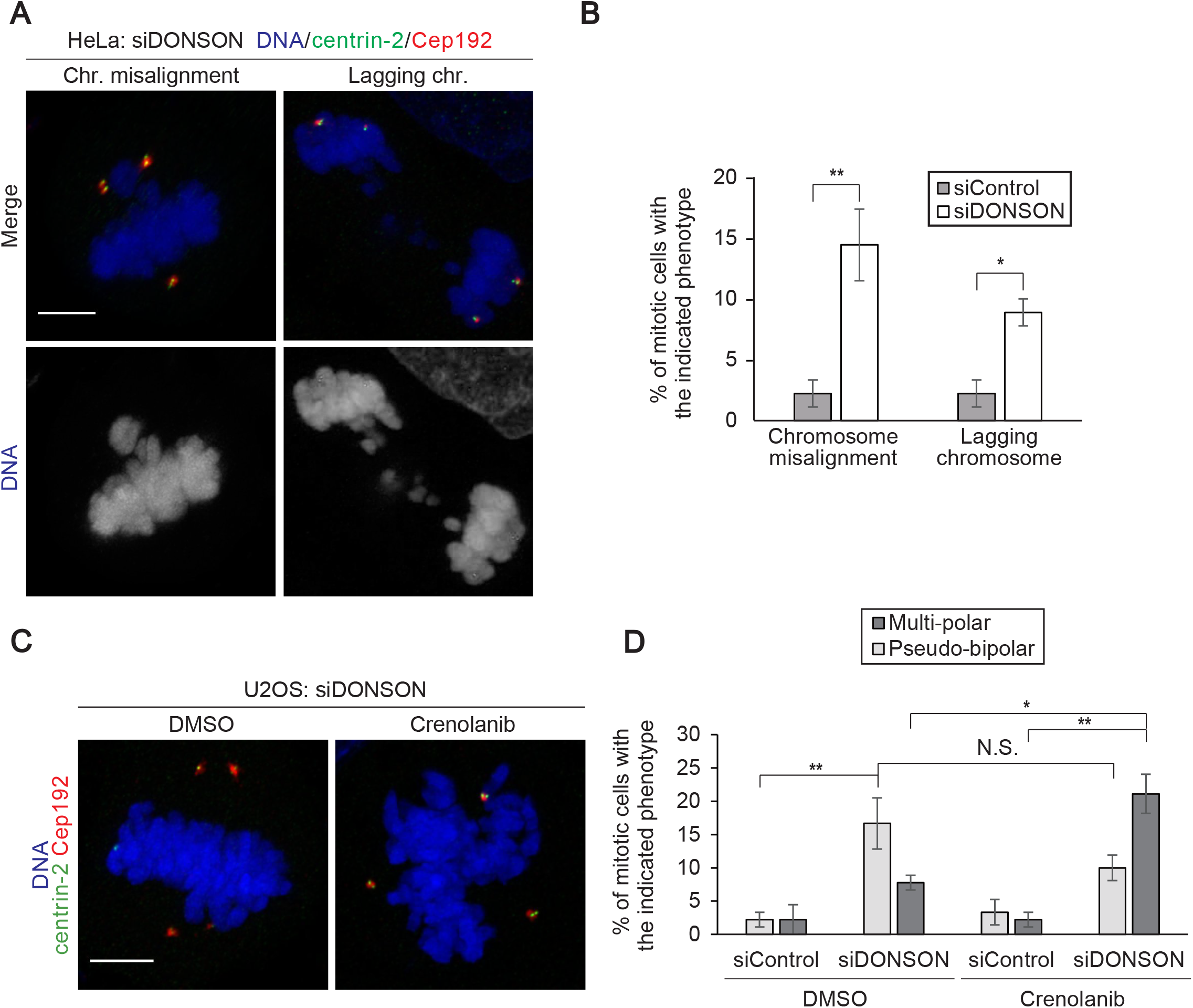
**A**, Chromosome segregation errors observed in DONSON-depleted HeLa cells. The cells were treated with siDONSON and immunostained with antibodies against centrin-2 (green) and Cep192 (red). **B,** Histograms represent frequency of mitotic cells with the indicated phenotype observed in A. Values are mean percentages ± s.d. from three independent experiments (N=30 for each experiment). **C,** Crenolanib treatment increased the number of mitotic cells with multi-polar spindles in DONSON-depleted U2OS cells. U2OS cells were treated with siRNA, followed by treatment with DMSO or Crenolanib (500 nM) for 6 h. The cells were immunostained as in A. **D,** Histograms represent frequency of mitotic cells with the indicated phenotypes observed in C. Values are mean percentages ± s.d. from three independent experiments (N=30 for each experiment). All scale bars, 5 μm. Two-tailed, unpaired Student’s t-test was used in B, and Tukey’s multiple comparison test was used in D to obtain P value. *, p < 0.05; **, p < 0.01; N.S., not significantly different (p > 0.05).

**Fig. S6.**
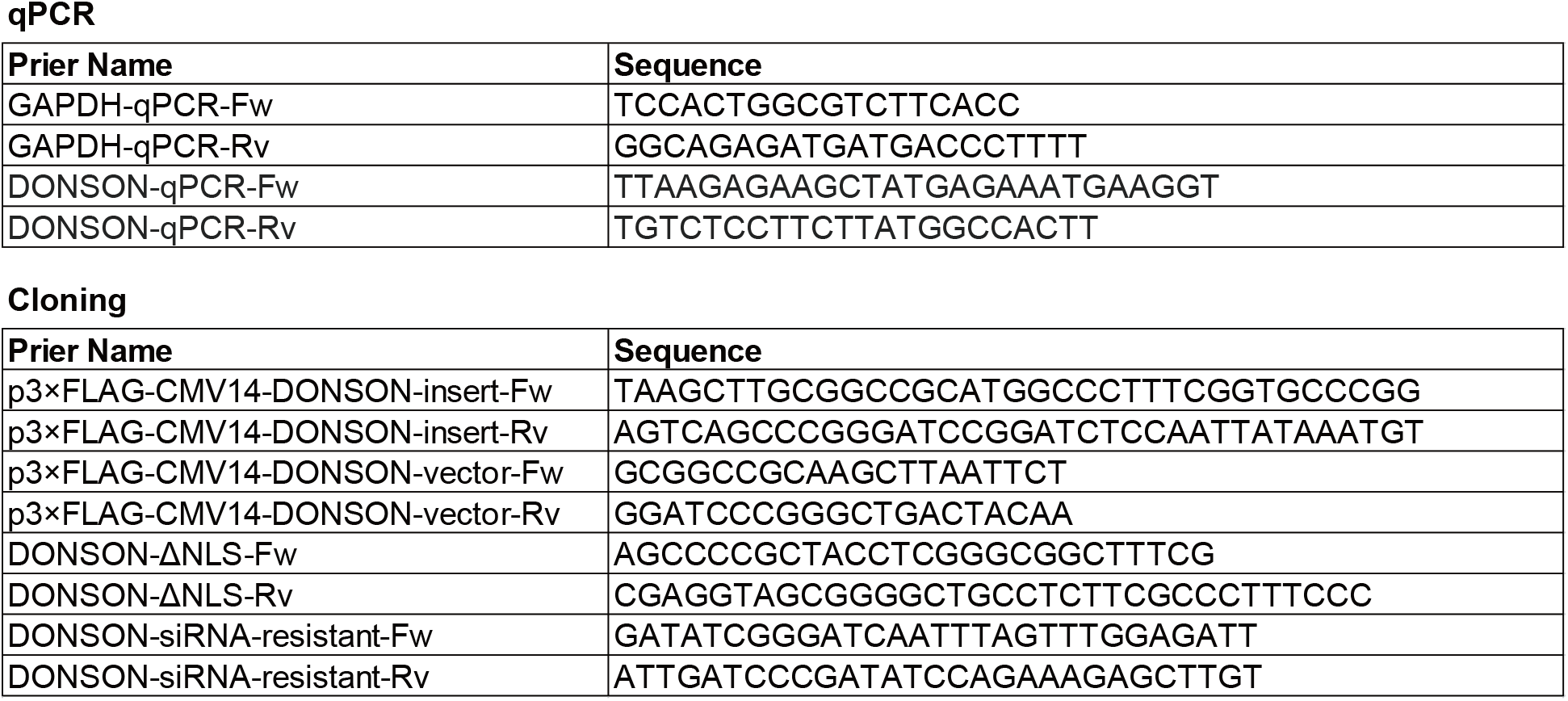
List of primers used in this study.

## Materials and Method

### Cell culture and transfection

HeLa, U2OS, RPE-1, and PANC-1 cells were obtained from the ECACC (European collection of cell cultures). HeLa Flp-In/T-Rex cells expressing GFP-DONSON, and U2OS cells were cultured in Dulbecco’s modified Eagle’s medium (DMEM), supplemented 10% fetal bovine serum (FBS) and 100 μg/ml penicillin–streptomycin at 37 °C in a 5% CO_2_ atmosphere. Growth media for HeLa Flp-In/T-Rex cells was additionally supplemented with 5 μg/ml blasticidin and 400 μg/ml hygromycin. 1 μg/ml doxycycline was added to the media for 24 h to induce expression of exogenous GFP-tagged DONSON. RPE-1 cells were cultured in DMEM / Ham’s F-12, supplemented 10% fetal bovine serum (FBS) and 100 μg/ml penicillin–streptomycin at 37 °C in a 5% CO_2_ atmosphere. PANC-1 cells were cultured in RPMI 1640, supplemented 10% fetal bovine serum (FBS) and 100 μg/ml penicillin–streptomycin at 37°C in a 5% CO_2_ atmosphere. hTERT immortalized fibroblasts derived from patients with *DONSON* mutations were infected with pMSCV-empty-vector or pMSCV-DONSON and were cultured in Dulbecco’s modified Eagle’s medium (DMEM), supplemented 10% fetal bovine serum (FBS) and 100 μg/ml penicillin– streptomycin at 37 °C in a 5% CO_2_ atmosphere. Transfection of siRNA or DNA constructs into HeLa, U2OS, and RPE-1 cells was conducted using Lipofectamine RNAiMAX (Life Technologies) or Lipofectamine 2000 (Life Technologies), respectively. Unless otherwise noted, the transfected cells were analyzed 48 h after transfection with siRNA and 24–36 h after transfection with DNA constructs.

### RNA interference

The following siRNAs were used: Silencer Select siRNA (Life Technologies) against DONSON #1 (s26833), DONSON #2 (s26834), Orc1 (s9893), Orc4 (s9898), Orc6 (s532063), Cdt1 (s532008), GMNN (s27306), Cdc6 (s2744), Cdc45 (s15830), Mcm5 (s8595), TP53 #1 (s529268), TP53 #2 (s529269), and negative control (4390843). Unless otherwise noted, DONSON #2 and TP53 #1 were used in this study.

### Plasmids

Complementary DNA (cDNA) encoding DONSON isoform (NCBI NP_060083.1) was amplified from cDNA library of HeLa cells. The DONSON cDNA was subcloned into p3×FLAG-CMV14 (Addgene). The DONSON ΔNLS mutant constructs were created using PrimeSTAR mutagenesis basal kit (TaKaRa) and In-Fusion HD cloning kit (Clontech) according to the manufacturer’s protocol.

### Drug treatment

The following chemicals were used in this study: Doxycyclin (Merck, D9891), Nocodazole (Wako, 31430-18-9), Crenolanib (Selleck, CP-868596), and SiR-DNA (Spirochrome, CY-SC007). For determination of S phase cells, cells were treated with 10 μM EdU for 30 min before fixation unless otherwise stated. The fixed cells were stained using Click-iT EdU Alexa Fluor 555 imaging kit (Life Technologies) according to the manufacturer’s recommendations.

### Antibodies

The following primary antibodies were used in this study: rabbit antibodies against Cep192 (Bethyl Laboratories, A302–324 A, IF 1:1000), CP110 (Proteintech, 12780-1-AP, IF 1:500), C-Nap1 (a gift from Erich Nigg IF 1:1000), CENP-F (Bethyl Laboratories, A301–617 A IF 1:1000), Cep152 (Bethyl Laboratories, A302–480A, IF 1:1000), PCNT (Abcam, ab4448, IF 1:2000), and γ-Tubulin (Merck, T5192, IF 1:1000),; mouse antibodies against centrin-2 (Merck, 20H5, IF 1:1000), HsSAS-6 (Santa Cruze Biotechnology, sc-81431, IF 1:1000), EB1 (BD Transduction Laboratories, 610534, IF 1:500), α-Tubulin (Merck, DM1A, IF 1:1000), GFP (Invitrogen, A-11120, IF 1:1000), and FLAG-tag (Merck, F1804, IF 1:1000). The following secondary antibodies were used: Alexa Fluor 488 goat anti-mouse IgG (H + L) (Molecular Probes, A-11001, 1:1000), Alexa Fluor 568 goat anti-rabbit IgG (H + L) (Molecular Probes, A-11011, 1:1000).

### Microscopy

For immunofluorescence analysis, the cells cultured on coverslips (Matsunami, 15 mm, thickness No. 1_0.13-0.17 mm) were fixed using −20 °C methanol for 7 min and washed with PBS. The cells were permeabilized after fixation with PBS/0.05% Triton X-100 (PBSX) for 5 min three times, and incubated for blocking in 1% BSA in PBSX for 20 min at room temperature (RT). The cells were incubated with primary antibodies for 24 h at 4 °C, washed with PBSX three times, and incubated with secondary antibodies for 1 h at RT. The cells were thereafter washed with PBSX twice, stained with 0.2 μg/ml Hoechst 33258 (DOJINDO) in PBS for 5 min at RT, washed again with PBSX, and mounted onto glass slides. Counting the number of immunofluorescence signals was preformed using an Axioplan2 fluorescence microscope (Carl Zeiss) with a ×63 or ×100/1.4 NA plan-APOCHROMAT objective and DeltaVision Personal DV-SoftWoRx system (Applied Precision) equipped with a CoolSNAP CH350 CCD camera. Confocal microscopy images were taken by the Leica TCS SP8 HSR system equipped with a Leica HCX PL APO ×63/1.4 oil CS2 objectives and excitation wavelength 405, 488 and 561 nm. To obtain high-resolution images, the pinhole was adjusted at 0.5 airy units. Scan speed was set to 200 Hz in combination with five-fold line average in 856 × 856 format (pixel size 43 nm). The images were collected at 130 nm z steps. For deconvolution, Huygens essential software (SVI; Scientific Volume Imaging) was used.

### Reverse transcription and real-time qPCR

Total RNA (1 μg) isolated from cells with the use of TRIZOL reagent (Thermo Fischer Scientific) was subject to reverse transcription using Quantiscript Reverse Transcription Kit (QIAGEN), and the resulting cDNA was subjected to real-time qPCR analysis with TB Green PCR Master Mix (TaKaRa) and specific primers in a StepOnePlus Real-Time PCR system (Applied Biosystems). The amount of each mRNA was normalized by that of GAPDH mRNA.

### Fluorescence activated cell sorting

Cells were trypsinized, washed twice with PBS, and fixed in ice cold 70% ethanol at −20°C for > 3 h. The fixed cells were then washed with PBS and incubated with Muse Cell Cycle Reagent for 30 min at RT in the dark. The DNA contents of the cells were then measured using Muse Cell Analyzer (Merck).

### Live imaging

Cell Voyager CQ1 (Yokogawa Electric Corp.) equipped with a ×40 oil immersion objective lens and the stage incubator for a 35-mm dish was used for live cell imaging. HeLa cells stably expressing GFP-centrin-1 or HeLa Flp-In/T-Rex cells expressing GFP-DONSON were cultured on 35 mm glass-bottom dishes (Greiner-bio-one, #627870) at 37 °C in a 5% CO_2_ atmosphere. Before imaging, cells were treated with siRNAs or 1 μg/mL Doxycyclin for 24 h and with 100 nM SiR-DNA for 3 h. Cells were visualized every 10 min over 24 h or 20 min over 48 h by a Back-illuminated sCMOS camera. The images were collected at 1.2 μm z steps. Maximum projections were generated using ImageJ (National Institutes of Health).

